# MotifScope: a multi-sample motif discovery and visualization tool for tandem repeats

**DOI:** 10.1101/2024.03.06.583591

**Authors:** Yaran Zhang, Marc Hulsman, Alex Salazar, Niccolò Tesi, Lydian Knoop, Sven van der Lee, Sanduni Wijesekera, Jana Krizova, Erik-Jan Kamsteeg, Henne Holstege

## Abstract

Tandem repeats (TRs) constitute a significant portion of the human genome, exhibiting high levels of polymorphism due to variations in size and motif composition. These variations have been associated with various neuropathological disorders, underscoring the clinical importance of TRs. Furthermore, the motif structure of these repeats can offer valuable insights into evolutionary dynamics and population structure. However, analysis of TRs has been hampered by the limitations of short-read sequencing technology, which lacks the ability to fully capture the complexity of these sequences. With long-read data becoming more accessible, there is now also a need for tools to explore and characterize these TRs. In this study, we introduce MotifScope, a novel algorithm for visualization of TRs in their population context based on a de novo k-mer approach for motif discovery. Comparative analysis against three established tools, uTR, TRF, and vamos, reveals that MotifScope can identify a greater number of motifs and more accurately represent the actual repeat sequence. Additionally, MotifScope enables comparison of sequencing reads within an individual and assemblies across different individuals, showing its applicability in diverse genomic contexts. We demonstrate potential applications of MotifScope in diverse fields, including population genetics, clinical settings, and forensic analyses.

## 1. Introduction

A large part of the human genome consists of repetitive elements. One such class of repeats are Tandem repeats (TRs), which are DNA sequences characterized by the contiguous repetition of at least one nucleotide, and they account for approximately 30% of the human genome (Audano et al. 2019; Gemmell 2021). TRs are broadly classified based on the size of their repetitive motif: TRs with motif size ≤ 6 bp are referred to as short tandem repeats (STRs), while those with larger motifs and variability in copy numbers are categorized as variable number tandem repeats (VNTRs) (Tautz 1993; Eslami Rasekh et al. 2021).

TRs are highly polymorphic, making them a major source of diversity in human genomes (Jeffreys et al. 1985). In fact, as few as 13 STRs in North America and 17 STRs in the United Kingdom, can uniquely identify a person (Hammond et al. 1994; Opel et al. 2007; Glynn 2022; Mallinder et al. 2022). Due to the high genetic variability and lack of linkage disequilibrium with each other, the probability of two unrelated individuals sharing a perfect match of this set of STRs is less than 1 in 1 billion (Reilly 2001). This variability has led to the adoption of these TRs as standard tools in forensics analyses, where they play a crucial role in DNA profiling and identifying individuals with a high degree of accuracy (Moretti et al. 2001; Jobling and Gill 2004).

Moreover, the variability observed in TRs extends beyond genetic diversity: the expansions of certain TRs have been associated with a range of neurological disorders (Chintalaphani et al. 2021; Hannan 2018; Tang et al. 2017). For instance, Friedreich ataxia (FRDA) can be caused by homozygous expansion of a “GAA” repeat in the first intron of frataxin (FXN) gene, while the expansion of a “GGGGCC” repeat intronic of C9orf72 gene has been linked to increased risk of amyotrophic lateral sclerosis (ALS) and frontotemporal dementia (FTD) (Pandolfo 2009; DeJesus-Hernandez et al. 2011). VNTR expansions have also been implicated in several diseases (De Roeck et al. 2018; Hannan 2018; Song et al. 2018). For example, the expansion of a 25 bp repeat in the intronic region of the ATP-Binding Cassette Subfamily A Member 7 (ABCA7) gene has been linked to an increased risk of Alzheimer’s disease (AD) (De Roeck et al. 2018). Interestingly, it is not just the repeat length that is associated with disease. In some cases, the motif composition of the repeat is also important (Chen et al. 2020; Cortese et al. 2019; Ishiura et al. 2018; Seixas et al. 2017; Wright et al. 2020). For instance, in patients with cerebellar ataxia with neuropathy and vestibular areflexia syndrome (CANVAS), expansions of the repeat in Replication Factor C Subunit 1 (RFC1) gene are composed primarily of “AAGGG” or “GACAG”, variations from the more common “AAAAG” motif found in non-expanded alleles (Cortese et al. 2019; Scriba et al. 2020).

Furthermore, because of their high mutability, TRs provide insight into evolutionary and population structure (Course et al. 2020; Ellegren 2004; Lu and Chaisson 2022; Rosenberg et al. 2002). Course et al. demonstrated significant differences in repeat length among VNTRs in genes including ART1, PROP1, and DYNC2I1, as well as substantial differences in motif organization in PCBP3 between superpopulations (Course et al. 2021). Using a pangenome-based approach, Lu et al. discovered that more than 8,000 VNTRs show differential motif usage across populations (Lu et al. 2021). These findings emphasize the importance of considering both repeat length and motif organization in TR genotyping.

With new sequencing technologies emerging that combine longer read length and higher accuracy, it is now possible to deeply characterize TRs. As such, multiple methods have been developed to genotype and profile TRs. One class of methods uses a provided sequence to detect motifs de novo. This allows them to handle extensive genomic diversity, including in complex TRs in which motifs can be highly variable. A drawback of these methods is that in a setting in which multiple individuals are analyzed together, it can be hard to canonicalize the discovered motifs across individuals. For instance, the widely used Tandem Repeats Finder (TRF) program (Benson 1999) generates a separate annotation for each motif it identifies, leaving it to the user to select the optimal motif representation, and to canonicalize these representations across different individuals. Recently, Masutani et al. proposed an algorithm, uTR, to decompose TRs after selecting a better set of motifs according to maximum parsimony that minimizes replication slippage events (Masutani et al. 2023). Still, this method will analyze each allele sequence individually. Furthermore, these methods are not sensible to small mutations (e.g., single nucleotide polymorphisms, SNPs). These small variations can however be biologically important, for instance in clinical settings in which pure repeats are considered more pathogenic than interrupted repeats.

Recently, another class of methods has been proposed which relies on databases of tandem repeats, describing for each TR the motifs that are to be considered. This has an advantage that discovered motifs are more homogenous across individuals. For instance, Ren et al. developed a toolkit, vamos, that generated a representative set of motifs for over 460,000 TRs in the human genome (Ren et al. 2023). This was achieved by selecting motifs for each TR in a reference set of genomes, while allowing for some sequence divergence. Then, vamos uses this motif database to annotate TRs in the query genome. Similarly, Dolzhenko et al. developed the Tandem Repeat Genotyping Tool (TRGT), specifically for PacBio HiFi sequencing data (Dolzhenko et al. 2024). It uses pre-specified motifs to genotype simple TR regions, while more complex repeats are defined using hidden Markov models (HMMs). Additionally, TRGT comes with a visualization module, the TRVZ tool, which displays read-level evidence supporting the genotype calls made by TRGT as well as the TR motifs. While the use of a motif database can work well for relatively stable TRs, it may result in the loss of important motifs for more variable TRs, for instance, rare motifs that are relevant to diseases. In addition, relying on a database hinders the application to novel repeats, or application to species not covered by the database (currently databases are only available for the human genome). Therefore, there remains a need for more versatile methods that can accurately capture the complexity of TR sequences across diverse genomic contexts.

Here, we present MotifScope, a flexible toolkit for motif annotation and visualization of TRs from sequencing data, that uses a de novo k-mer-based approach for motif discovery. To evaluate MotifScope performances, we compared it to the three existing tools for motif discovery: uTR, TRF and vamos. Our findings indicate that MotifScope identified a greater number of motifs and reflected the actual repeat sequence more accurately than other tools. Additionally, we show potential applications of MotifScope in population genetics to explore population stratification due to TRs, in clinical studies to study pathogenic TRs, and in a forensic setting.

## 2. Result

### 2.1. Accuracy and efficiency

To evaluate the performance of MotiScope in characterizing TRs, we conducted a comparative analysis alongside recently developed TR-analysis methods: TRF, uTR, and vamos (with both the original motif set, “vamos original”, and an efficient motif set, “vamos efficient”) on the HG002 genome. As for TRs set, we used the set of 5,486 repeats that are present in both the PacBio repeat catalog and the vamos VNTR catalog.

MotifScope identified a greater number of motifs compared to the other three tools (Figure 1A). To evaluate the quality of motif identification, we assessed the edit distance between the concatenation of the motif representations generated by these tools and the true repeat sequences (Figure 1B). By design, MotifScope consistently achieves an edit distance of 0, indicating an exact match between its motif description and the underlying repeat sequence. In contrast, other methods exhibited increasing edit distances, for example, by sorting the edit distance in ascending order, at 90th percentile, the edit distance was 0 for MotifScope, 0.022 for TRF, 0.032 for vamos original, 0.029 for vamos efficient and 0.098 for uTR. This shows that MotifScope’s description of TRs reflects the actual repeat sequences more accurately compared to the other three tools.

**Figure 1.**
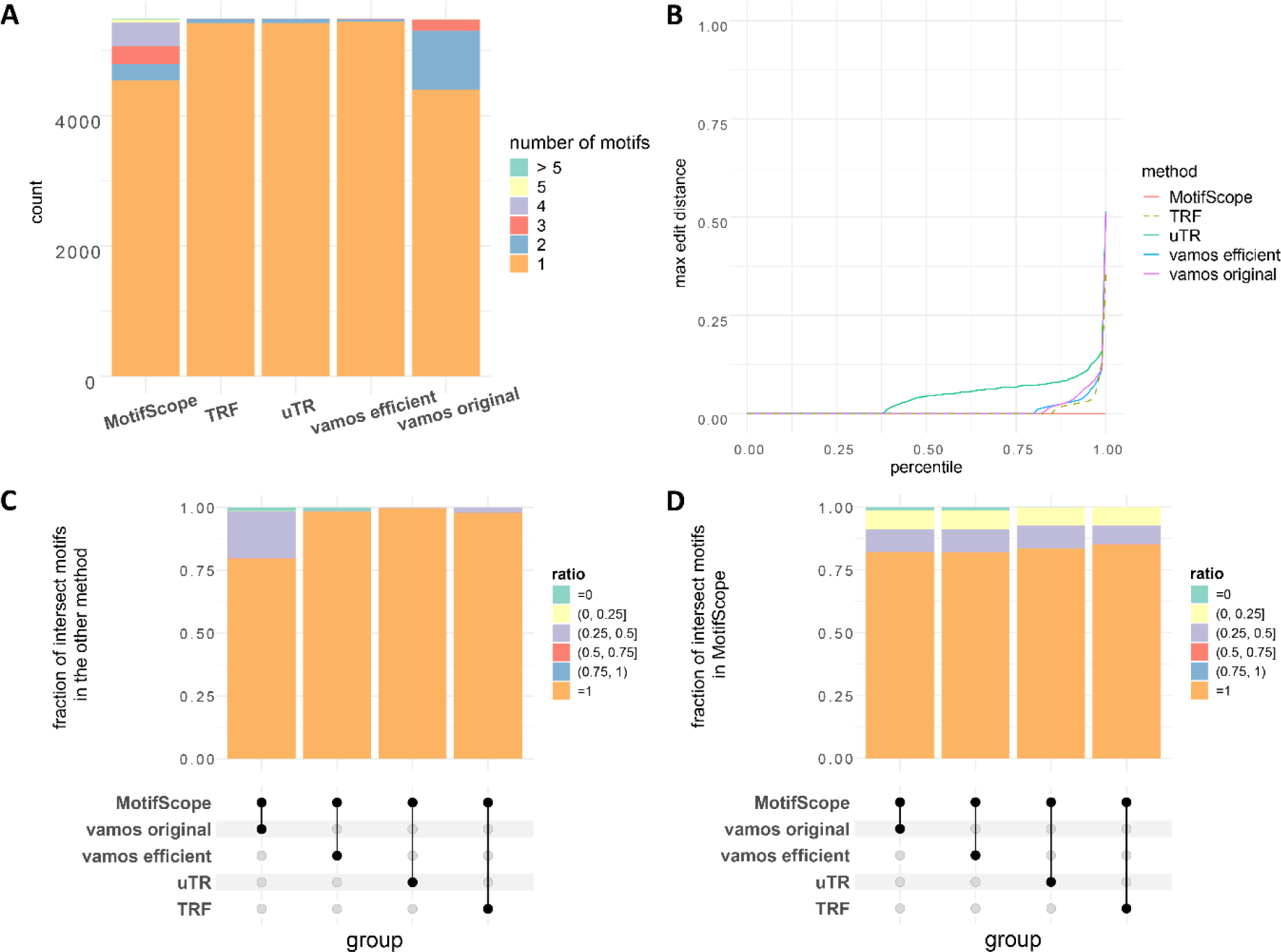
Comparative analysis of tandem repeat characterization. Four tools were tested in the analysis, MotifScope, TRF, uTR, and vamos. For vamos, here we used both the original motif set (vamos original) and the efficient motif set (vamos efficient). (A) shows the number of motifs discovered by each of the four tools. (B) shows the normalized edit distance between the actual sequence and the results obtained from the four tools, normalized with respect to repeat length. For MotifScope, uTR and vamos, the normalized edit distance is calculated using the concatenation of the motif representation of the repeat with the entire actual sequence of the repeat. In the case of TRF, where multiple results were sometimes provided for a single repeat, the normalized edit distance is calculated using the results from TRF (i.e., motif * copy number) and the true underlying sequence of that explanation. The average of these values was used to represent the normalized edit distance for TRF. (C) and (D) shows the intersection of motifs between MotifScope and other tools (dots connected by lines below the X axis): in the bar plot in the middle, each column shows the number of loci with results obtained from both MotifScope and the respective other tools; for (C), the stacked bar plot at the top shows the fraction of intersected motifs over the total number of motifs found by the other tool; for (D) the stacked bar plot at the top shows the fraction of intersected motifs over the total number of motifs found by MotifScope.

We further evaluated the extent to which different tools captured the same motifs within the studied TRs: we found that MotifScope identified all motifs found by TRF in 97.89% of loci, by uTR in 99.75%, by vamos original in 79.64%, and by vamos efficient in 98.34% of the loci (Figure 1C). The relatively lower discovery rate of motifs from vamos original by MotifScope is attributed to the presence of rare motifs (occurring only once) in vamos original motif sets, which are characterized differently by MotifScope as single nucleotides. Moreover, MotifScope was also able to pick up motifs that were not detected by the other tools. For example, in 15.30% of TRs, MotifScope found motifs not identified by TRF, in 16.49% not identified by uTR, in 17.70% and 17.94% not identified by vamos original and vamos efficient, respectively (Figure 1D). These additional motifs, including single nucleotides found by MotifScope, allow Motifscope to more accurately represent the underlying repeat sequence. In the case of the 5,000 vamos VNTR, MotifScope also identified more motifs and reflected the sequence more accurately compared to the other tools, however, the overlap of motifs identified by MotifScope and other tools decreased in this subset, reflecting the high complexity of this set of repeats (Supplementary Figure 1).

### 2.2. Merit of de novo motif discovery

MotifScope uses a de novo motif discovery approach to enable the discovery of rare, possibly pathogenic motifs. In Figure 2, we present the motif characterization results of a TR within intron 2 of RFC1 (chr4:39348424-39348483), where biallelic AAGGG or GACAG repeat expansion has been associated with an autosomal recessive neurological disorder, cerebellar ataxia with neuropathy and vestibular areflexia syndrome (CANVAS). We employed MotifScope to characterize this repeat on HG002 and a Dutch CANVAS patient (Wang et al. 2022; van de Pol et al. 2023). The patient is a carrier of homozygous expansions of 6.28 and 7.69 kb for the 2 haplotypes. In HG002, all four tools annotated the repeat using the motif AAAAG or its cyclic shifts. In the CANVAS patient, MotifScope identified three major motifs: GACAG, GACAA, and AAAAG. Using these motifs, the TR can be characterized as (GACAG)_1136_(GACAA)_118_GAAG(AAAAG)_14_ with 4 indels for haplotype 1 and (GACAG)_1420_(GACAA)_114_GAAG(AAAAG)_15_ with 20 indels for haplotype 2. Note that the results of MotifScope also represent the indels through additional motifs (total of 6 motifs), resulting in a fully accurate representation of the actual repeat sequences (edit distance = 0.0). In contrast, uTR annotated the sequences using GACAG and GACAA, (GACAG)1137(GACAA)133 for haplotype 1 and (GACAG)1422(GACAA)130 for haplotype 2, with an average edit distance of 0.005, mislabeling the AAAAG motifs at the end the alleles. TRF explained the sequences with a single GACAG motif, (GACAG)1255 for haplotype 1 and (GACAG)1535 for haplotype 2, resulting in an average edit distance of 0.019. Finally, vamos original primarily characterized the sequence with motifs “GGGAC” and “AAAAG”, (GGGAC)1177(AAAAG)130 for haplotype 1 and (GGGAC)1418(AAAAG)130 for haplotype 2 generating an average edit distance of 0.212, while vamos efficient predominantly used motifs “GGAAA”, “GGCAA”, and “AAAAG” for annotation, resulting in (GGAAA)1181(GGCAA)115(AAAAG)14 for haplotype 1 and (GGAAA)1422(GGCAA)115(AAAAG)14 for haplotype 2, with an average edit distance of 0.214.

**Figure 2.**
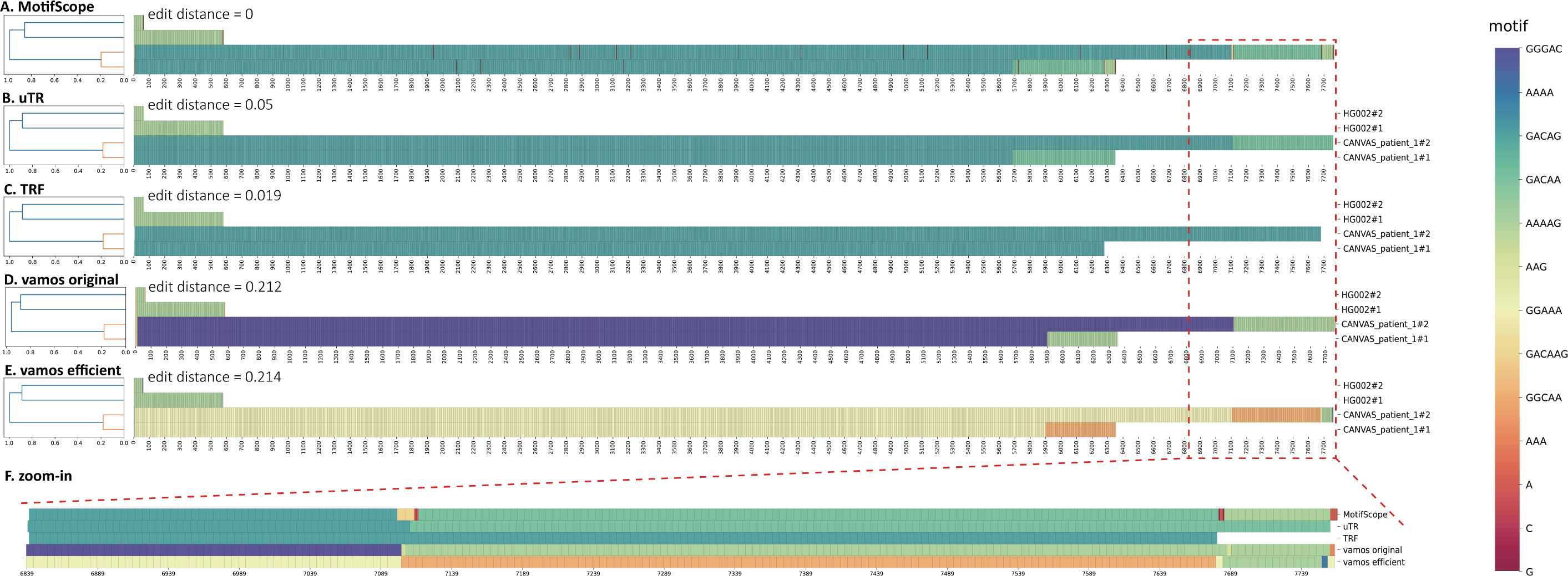
Motif characterization of the RFC1 repeat in HG002 and a CANVAS patient. The results from (A) MotifScope, (B) uTR, (C) TRF, (D) vamos original, (E) vamos efficient of the RFC1 repeat in the HG002 assembly and the assembly of a Dutch CANVAS patient, and (F) zoomed-in view of the circled region of one patient allele are visualized. In each subfigure, the left panel shows the clustering of the sequences, the right panel shows the composition of the repeat, with distinct motifs represented in different colors, as indicated by the color bar on the right side of the figure.

To further illustrate the motif discovery results, we also present the characterization of a forensic locus, D2S441 (chr2:68011946-68011994), in the HG002 genome assembly (Figure 3). MotifScope and vamos original successfully characterized the two alleles as (TCTA)_11_ and (TCTA)_12_TTTA(TCTA)_2_ while uTR, TRF and vamos efficient failed to identify the “TTTA” motif in the second allele.

**Figure 3.**
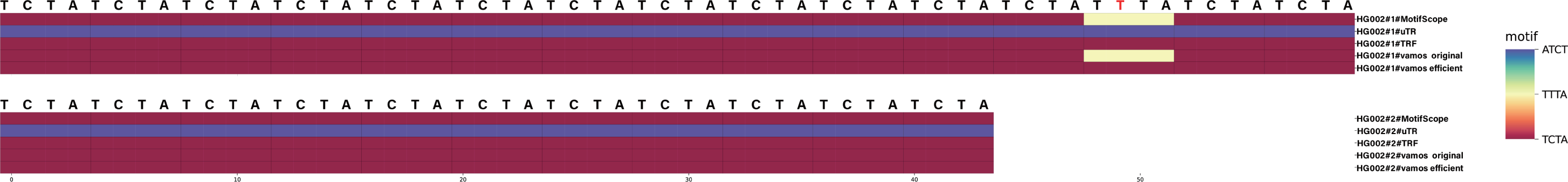
Motif characterization of forensic loci D2S441 in HG002. This shows the visualization of the decomposing result from MotifScope, TRF, uTR, vamos efficient and vamos original of the local assembly of the forensic D2S441 locus in HG002 genome assembly, with distinct motifs represented in different colors, as indicated by the color bar on the right side of the figure.

### 2.3. Joint motif analysis across individuals

MotifScope offers the capability to analyze multiple samples simultaneously, which can improve the characterization and comparison of TR haplotypes. For example, when MotifScope was applied to the RFC1 repeat in a single genome HG01175 from the Human Pangenome Reference Consortium (HPRC), it initially failed to identify the starting motif “AAAAG” and motif “GACGG” immediately after the “AAAGG” repeat in haplotype 1. Instead, these motifs were represented as single nucleotides (e.g., “A”, “A”, “A”, “A”, “G” for “AAAAG”) (highlighted in the red box in Figure 4A). As the “GACGG” and “AAAAG” motifs are in the original motif set, vamos original was able to identify these two single motifs on HG01175 (Supplementary Figure 2). However, when jointly analyzing HG001175 with HG01109 and HG00733, MotifScope managed to also identify the single copies of these motifs in HG001175, as these were recurring motifs across samples. In addition, joint analysis also revealed that HG01175 haplotype 1 and HG01109 haplotype 1 share the same motif structure: AAAAG(AAGGG)_7_ (GACGG)_1/2_(AAAGGG)_n_(AAAGGGAAGG)_2_AAAG(GAAA)_2_AAG (Figure 4D).

**Figure 4.**
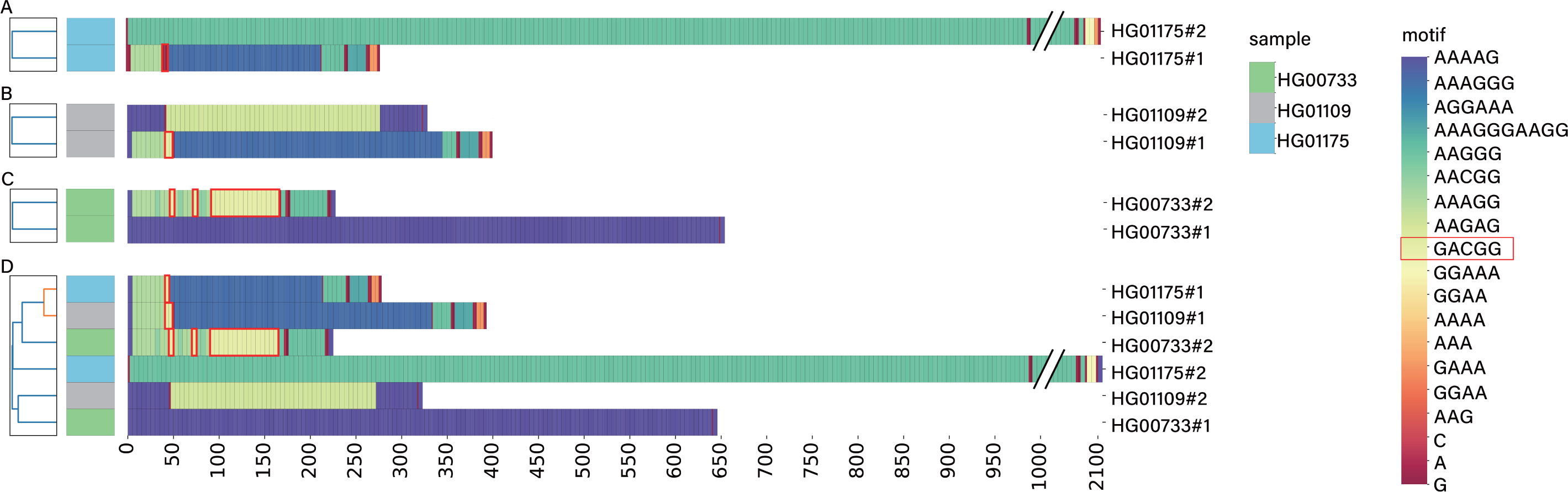
Motif characterization of the RFC1 repeat in three HPRC genomes. The results from MotifScope on the assemblies of three genomes, (A) HG01175, (B) HG01109 and (C) HG00733. These three genomes are Admixed Americans. (D) shows the joint analysis result of these three genomes. The sequence “GACGG” in these sequences are highlighted in red boxes. The left panel shows the clustering of sequences along with genome identifiers, represented by the corresponding color bar on the second-to-right side. The right panel shows the motif composition of the repeat, with distinct motifs represented in different colors, as indicated by the color bar on the right side of the figure.

### 2.4. Profiling clinically relevant loci in the population

TRs are known to have a wide variability in motif sequence, motif size, and repeat size across populations, suggesting the importance of examining multiple samples from different populations collectively. For example, a pathogenic repeat in BEAN1 (chr16:66490397-66490466) is associated with an autosomal dominant neurological disorder, Spinocerebellar Ataxia Type 31 (SCA31), and coincides with a specific “GAATG” repeat insertion found solely in the Japanese population (Ishikawa and Nagai 2019). We applied MotifScope to this locus using 47 genome assemblies from the HPRC, and identified a cluster of African alleles with distinct motif structure, with “A” and “T” insertions in the AAATA repeat (in purple box in Figure 5). Additionally, two expanded alleles were observed: a 2.47 kb expansion with “CAATA” motifs in an admixed American allele and a 0.99 kb “AAATA” expansion in an East Asian allele. Notably, we observed expansions with “CAATA” and “AAATA” motifs as well as another motif “CACTA” that was found in a South Asian individual (Ishikawa and Nagai 2019).

**Figure 5.**
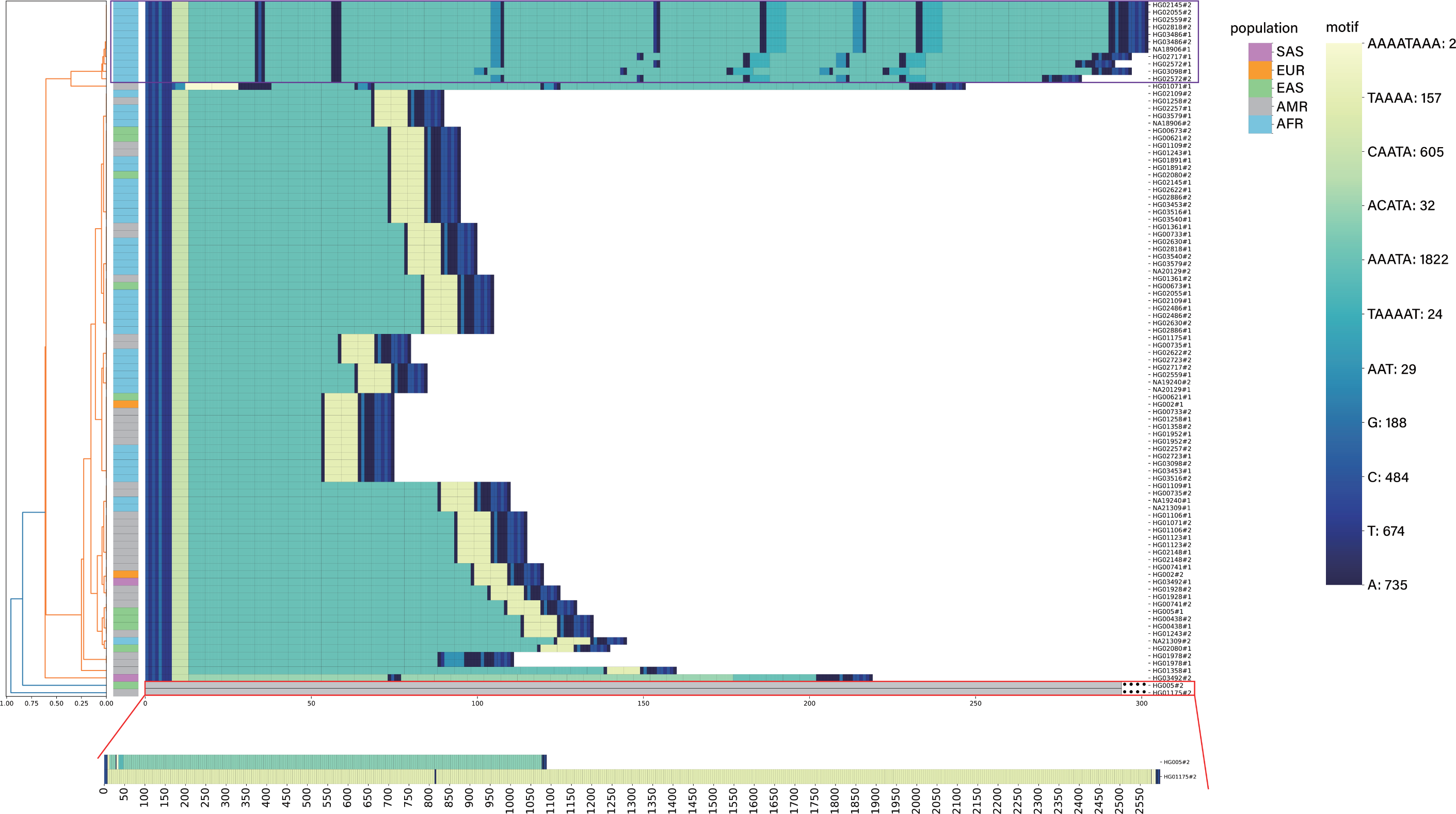
BEAN1 repeat in HPRC samples. The leftmost panel presents the clustering of assembly sequences from HPRC sample (n = 47) with 10 bp flanking the repeat, along with population origin of the alleles in the adjacent column. The color code, denoting population origin, is in the second-to-right color bar (SAS: South Asian; EUR: European; EAS: East Asian; AMR: Admixed American; AFR: African). The rightmost panel visually represents repeat composition, with distinct colors signifying different motifs. The sequences contain 10 bp flanking sequences on both sides of this locus. The corresponding color bar is on the right side of the figure. The number following the motif on the color bar indicates the number of occurrences of the motif in the figure. The two expanded alleles are truncated in this figure. The motif compositions of these two alleles are shown in (B).

We also applied MotifScope to a pathogenic repeat recently identified in FGF14 (chr13:102161575- 102161726) (Figure 6) (Rafehi et al. 2023; Pellerin et al. 2023). We used the assemblies of the HPRC samples as well as 250 Dutch AD patients and 238 Dutch centenarians. The expansion of this repeat has been associated with an autosomal dominant adult-onset ataxia SCA27B. Despite sharing the same “AGA” repeat backbone, 10 different repeat structures were identified across all evaluated individuals. Similar to the GRCh38 human reference genome, the majority of the alleles carried a non-expanded “AGA” repeat. However, the other assemblies carried different insertions inside the “AGA” repeat; insertions of single “A”s, an “AAGAGG” repeat insertion, an “AGG” repeat insertion immediately followed by a “GAGAAG” repeat insertion, an [AG(AGG)_2_A] repeat insertion, a “GAGAAG” repeat insertion, insertions of different single nucleotides in slightly expanded “AGA” repeat, and a East Asian individual with an 1.85 kb expansion composed of an [AGC(AGA)_4_] repeat, an “AAGCAG” repeat, and an “AGC” repeat insertions (Figure 6). Notably, MotifScope revealed that all African individuals had the same starting sequence of the repeat as the non-expanded “AGA” alleles whereas individuals of other ancestry that did not carry the pure non-expanded “AGA” allele had a different starting sequence (Figure 6). This case illustrates that the motifs and motif structures of TRs can be highly variable in the population and MotifScope is able to identify them and cluster sequences with the same motif structure together.

**Figure 6.**
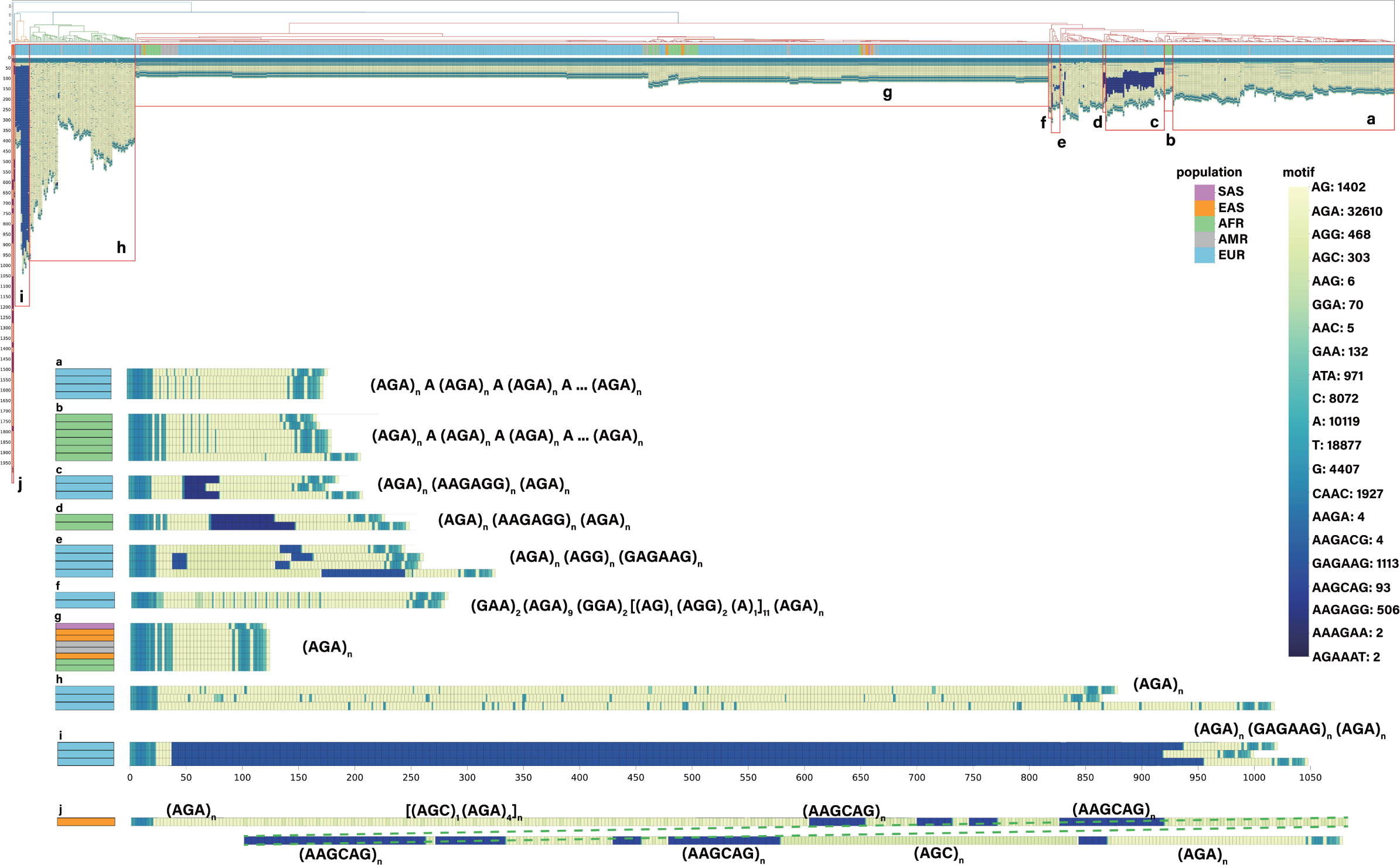
FGF14 repeat in HPRC, Dutch AD patients and centenarians. The assembly sequences on the FGF14 repeat with 31 bp flanking the repeat, of 47 HPRC samples, 250 Dutch AD patients and 238 Dutch centenarians. The top figure showed the result from MotifScope. The alleles below showed the zoomed in of different alleles that are present in these assemblies with the left column indicating the corresponding population (second-to-right color bar, SAS: South Asian; EUR: European; EAS: East Asian; AMR: Admixed American; AFR: African) of the assembly and the right figure showing the motif structure of these different alleles. The corresponding motif color bar is on the right side of the figure. The number following the motif on the color bar indicates the number of occurrences of the motif in the figure.

We also applied MotifScope to an intronic repeat in gene ABCA7 (chr19:1049437-1050066) on HG002, where the motif size is 25 bp (Supplementary Figure 3). MotifScope was able to identify 24 different motifs in this sequence including the 4 single nucleotides. However, a substantial portion of the sequence was annotated with these single nucleotides, posing a challenge for interpretation.

### 2.5. Analyzing TRs at read-level

Whereas MotifScope can jointly analyze multiple individuals, it can also jointly characterize and visualize all mapped sequencing reads that span a TR region, which can unveil somatic and technical variation. For instance, in Figure 7, we display all spanning reads from the blood of a Dutch CANVAS patient on the RFC1 repeat, revealing a motif structure characterized by (GACAG)_n_(GACAA)_n_(AAAAG)_n_. However, these reads varied in size, ranging from 7.0 kb to 9.5 kb, and contained different SNVs at different positions.

**Figure 7.**
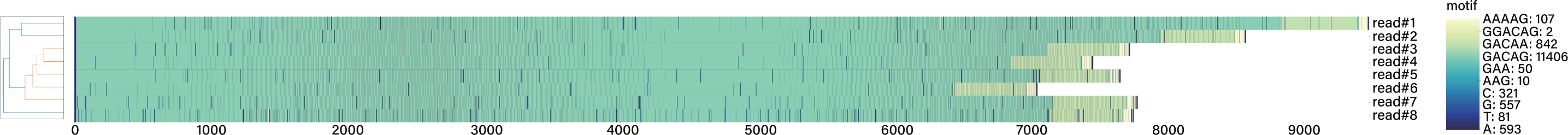
Motif characterization of the spanning reads of the RFC1 repeat in the blood of a CANVAS with MotifScope. The sequences contain the RFC1 repeat and 10 bp sequences flanking both sides of this region. The clustering of the sequences is shown in the left panel, the right panel shows the motif composition of the repeat, with distinct motifs represented in different colors, as indicated by the color bar on the right side of the figure. The number following the motif on the color bar indicates the number of occurrences of the motif in the figure.

Another example is shown in Figure 8, where we analyzed all reads as well as the phased assemblies for the forensic locus D3S1358 (chr3:45540738-45540802) in HG002 together using the multiple sequence alignment (MSA) feature provided by MotifScope. Our analysis revealed that 14 out of 31 reads matched one assembly allele, (TCTA)_1_(TCTG)_2_(TCTA)_12_, 12 out of 31 reads matched the other assembled allele, (TCTA)_1_(TCTG)_1_(TCTA)_14_, while 5 out of 31 reads did not exactly match to any of the two assembly alleles. Of these 5 reads, 3 reads contained a whole motif gain or loss compared to the two assemblies, 1 read contained a single “C” deletion, and 1 read contained a “C” insertion in addition to a whole “TCTA” motif loss. This analysis allows one to assess somatic stability and/or the propensity for technical sequencing errors in a region.

**Figure 8.**
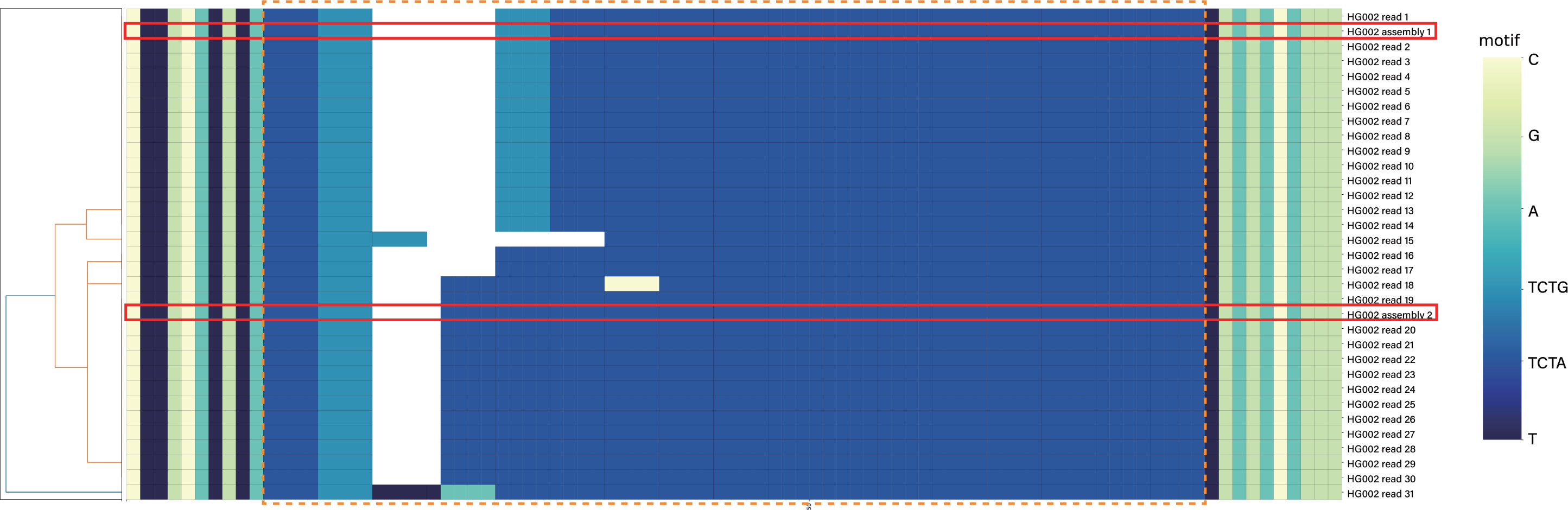
Motif characterization of the spanning reads of forensic loci D3S135 with MotifScope. The two assembly alleles of the HG002 genome were highlighted in red boxes and the repeat is highlighted in the orange box with 10 bp flanking this region. The clustering of the sequences is shown in the left panel, the right panel shows the motif composition of the repeat, with distinct motifs represented in different colors, as indicated by the color bar on the right side of the figure.

## 3. Discussion

Advancements in long read sequencing technologies have significantly enhanced our ability to explore TRs within the human genome. These variants have emerged as crucial elements in genetic studies due to their implications in evolution and diseases (Perry et al. 2008; Weischenfeldt et al. 2013). They often display multiallelic patterns in the human population, and therefore, a detailed examination of the composition of these repeats is important for unraveling their functional implications and evolutionary history. To address the complexities of TRs, we developed MotifScope, a tool designed to characterize and visually represent TR compositions. In a benchmarking study against existing tools, TRF, vamos, and uTR, MotifScope outperformed by identifying more motifs and more accurately reflecting sequence composition.

By performing motif discovery de novo, MotifScope does not have to rely on a predetermined set of motifs for TR decomposition, which makes it able to detect rare or unseen motifs. This is especially important in a clinical setting, as studies have shown that motif alterations in repeats can be the cause of disease. For instance, in CANVAS patients, pathogenic alleles of the RFC1 repeat have been identified as expansions of “GACAG” and “AAGGG” sequences, rather than the “AAAAG” repeat observed in the GRCh38 human reference genome (Cortese et al. 2019; Scriba et al. 2020; van de Pol et al. 2023). Similarly, in five familial adult myoclonic epilepsy (FAME) subtypes, expanded “TTTCA” segments were found inside the “TTTTA” repeat in SAMD12, STARD7, MARCHF6, TNRC6A and RAPGEF2 genes (Bennett et al. 2020; Corbett et al. 2019; Florian et al. 2019; Ishiura et al. 2018). Given that both the original and efficient set of motifs from vamos were constructed with the HPRC and Human Structural Variation Consortium (HGSVC) samples, some of these pathogenic motifs are not represented in the default motif sets (Figure 2) (Ren et al. 2023). Consequently, there is a risk of overlooking these rare, biologically relevant motifs, which could have significant implications for disease diagnosis and understanding.

MotifScope also offers the capability for joint motif analysis across samples. Similar to vamos, which utilizes motifs present in the HPRC and HGSVC samples, MotifScope thereby uses sequence information from other samples to enrich motif characterization (Figure 4). This allows it to correctly annotate single occurrences of a motif in a haplotype, by leveraging information from other genomes in which the motif is expanded. Such patterns can highlight patterns that reflect evolutionary proximity. Furthermore, joint analysis in combination with motif canonicalization ensures consistent annotation of motifs across all sequences. This prevents the occurrence of different motifs and/or cyclic shifted forms of the same motifs in different sequences, enhancing comparison across sequences.

Such comparisons will also highlight population differences. Several studies have demonstrated that while overall patterns of repeat variations are highly similar across populations, there are notable exceptions with population-specific patterns (Ziaei Jam et al. 2023). For instance, common “CAG” expansions in an intronic repeat within the gene CA10 were found predominantly in African individuals, and the motif usage of an intronic repeat in gene PCBP3 showed substantial differences across modern superpopulations (Ziaei Jam et al. 2023; Course et al. 2021). Another example is the BEAN1 repeat, which showed population-specific pathogenic expansions (Figure 5) (Ishikawa and Nagai 2019). Ataxia SCA31, found exclusively in the Japanese population, aligns with a specific “GAATG” repeat insertion in this repeat, which is also exclusive to the Japanese population, suggesting a strong founder effect. This insertion ranges 2.5–3.8 kb in size and was found to inversely correlate with disease age of onset. Our analysis of the FGF14 repeat in the HPRC samples, Dutch AD patients, and Dutch cognitively healthy centenarians (Figure 6) also revealed highly population specific patterns and revealed that only certain haplotype clusters showed expansions.

Repeat sequences might also differ at the read level. Somatic instabilities within TRs have been widely observed, including in forensic STRs (Figure 8) and pathological conditions such as cancers and repeat expansion disorders like Huntington’s disease, myotonic dystrophy type 1 and CANVAS (Figure 7), contributing to variability at the individual read level (dos Santos et al. 2012; Monckton 2021; Veitch et al. 2007; Ciosi et al. 2019; Chintalaphani et al. 2021). Simultaneously, despite the high accuracy of long read sequencing technologies, such as PacBio HiFi sequencing with an error rate lower than 0.05% and Oxford Nanopore with R10.4 with an error rate lower than 0.09%, errors can still occur during sequencing, leading to variations between individual reads. This can make it challenging to separate somatic from technical variation. However, we find that variations in reads also include whole-motif gain or loss in addition to SNVs. Given that homopolymer errors (i.e., indels) are the primary source of long-read sequencing systematic errors when using the PacBio technology, whole-motif gain and loss might be reflective of somatic variations (Au et al. 2012). With MotifScope, we can examine and separate these different classes of variations between reads while annotating them with a consistent motif set.

Alignment of sequences can further facilitate repeat sequence comparisons. However, traditional alignment methods may struggle to accurately align highly repetitive sequences, in which motif gains and losses occur, while only subtle differences can be used as alignment markers. Standard alignment approaches therefore do not always yield the most meaningful results. Here, we addressed this issue by aligning sequences based on their motif composition, performing alignment on so-called “motif sequences”, which lead to a more biologically relevant alignment. This approach also enhances the visual representation and comparability of repeat loci, particularly for analyzing differences between reads (Figure 8) in complex loci and identifying variations among haplotypes with similar motif structures.

One element which sets MotifScope apart is that motifs are annotated exactly as they appear in the input sequences, providing a precise representation of repeat structures. In contrast, other tools usually allow for some flexibility in describing repeat sequences. This leads to simpler more condensed motif patterns, but can also hide crucial information. In particular, small variations can highlight evolutionary proximity and expansion locations. Moreover, small variations can also be highly relevant in clinical settings, in which certain repeats have been found to only have pathogenic effects when the repeat is pure, i.e. is not interrupted by small sequence variations. One example is an intronic repeat in gene FGF14. Expansion of this repeat (> 250 repeats) can cause adult-onset ataxia SCA50/ATX- FGF14 (Rafehi et al. 2023). Previous studies have found that only those with pure “GAA” repeat expansions develop the disease and it was hypothesized that it is because these long “GAA” repeats are known to form secondary structures and therefore inhibit the transcription of the gene (Rafehi et al. 2023). With MotifScope, the structure of the repeat can be easily checked and visualized. As shown in Figure 6, two individuals had > 250 “GAA” repeats, yet they all have SNVs within the repeat sequence, and none were given ataxia diagnosis. However, it’s important to note that these samples have low coverage; two of these three haplotypes had two supporting reads and one had one supporting read, suggesting the possibility of sequencing errors affecting both the purity and size of the repeats.

Due to biological variations in TRs and technical errors introduced during sequencing, TR sequences often deviate from a perfect repetition of a single motif. Despite these variations, k-mers that form longer continuous sequence stretches are still more likely to explain more of the sequences and are consequently the frequently observed motifs. While this approach is effective for active repeats in which motif copies have not substantially diverged, it can also present a limitation in analyzing highly diverged sequences (Supplementary Figure 1). Exact motif discovery approaches can struggle with sequences in which repeats are hard to identify due to the buildup of mutations. This is more likely to occur in VNTRs with large motifs. For instance, Supplementary Figure 3 shows an example of an intronic ABCA7 repeat, where the sequence is highly complex. While the concatenation of results from MotifScope produces the true sequence, these additional motifs do not enhance the interpretation of the repeat’s structure, as much of the sequence is labeled with single nucleotides. MotifScope prioritizes motifs that can annotate long consecutive sequences, but when nearby motifs are all different, it fails to identify them. In such cases, tools like uTR, which allow distances to the true sequence, or vamos, which utilizes an established motif set to decompose repeat sequences, may be more suitable alternatives. Future iterations of MotifScope might therefore benefit from incorporating a more robust initial repeat structure identification step. Exact motif descriptions of highly complex repeats in combination with the here proposed motif color mapping could thereby reveal both the high-level structure of the repeat locus as well as more recent patterns of motif divergence.

With the continued adoption of long-read data in research and clinical settings, characterization and visualization of tandem repeats will remain an active research area, as TRs constitute a major source of biological variation (Jeffreys et al. 1985). MotifScope offers a new unique angle to this, which might lead to new insights into motif structure and pathogenic mechanisms. Joint motif analysis thereby facilitates sequence comparisons, which is leveraged in assembly-pileups for comparative analyses of motif structures across individuals, as well as read-pileups for analyzing somatic differences within an individual. The joint analysis of tandem repeat motif structures might also open up new avenues for case/control studies to not only take into account repeat length but also repeat structure in a biological meaningful manner. This could shed new light on the biological and phenotypical impact of tandem repeats.

## 4. Methods

MotifScope aims to characterize and visualize motif organization of TRs. It is designed to specifically target TRs that are variable among individuals. Although it works best with genome assemblies, it can be applied to any collection of genomic sequences (e.g., individual long-reads) from different technologies (e.g., PacBio, Nanopore).

MotifScope accepts a .fasta file containing tandem repeat sequences as input. It can be run in three modes: *assembly* mode for assessing variations between individuals, *reads* mode for comparing (somatic) variations within sequencing reads obtained from a single individual, and *single-sequence* mode for analyzing the motif structure of one sequence. Optionally, motif discovery can be guided by providing an expected set of motifs (in a .tsv file). Finally, MotifScope can be used for performing multiple sequence alignment (MSA) based on motif composition. MotifScope outputs the motif composition in a fasta file (including each motif and the relative number of copies) and a visual representation of motif composition.

MotifScope has been written in Python (version>=3.10). The code, documentation and example files are publicly available at https://github.com/holstegelab/MotifScope.

There are four major algorithmic steps in MotifScope: (1) iteratively identifying and annotating motifs using a k-mer-based approach; (2) color mapping of motifs using a dimensional reduction technique, (3) clustering and aligning sequences based on their motif composition; and (4) automatically generating a set of figures to display motif composition and similarity across sequences (Figure 9). MotifScope prioritizes motifs that make up long continuous sequences and therefore describes the organization of the frequently repeated motifs in TRs.

**Figure 9.**
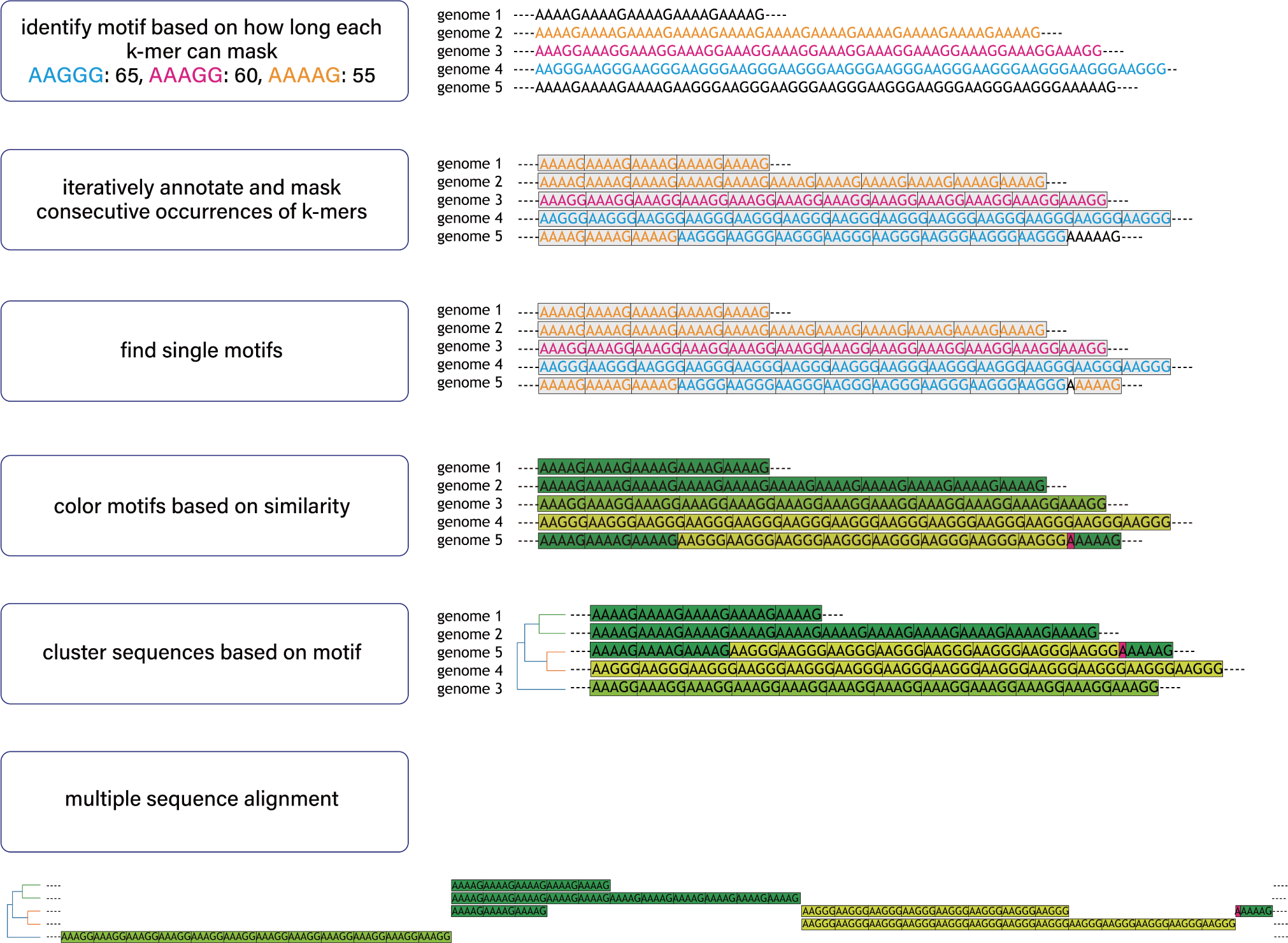
Overview of MotifScope. Input to MotifScope consists of a set of sequences. MotifScope first identifies motifs by evaluating the length of consecutive sequence stretches formed by each k-mer present in the sequences. It then iteratively annotates consecutive occurrences of k-mers as motifs and masks them in the sequences. Subsequently, single occurrences of identified motifs are annotated and masked. To visualize the motif composition, motifs are colored based on their sequence similarity to each other. The sequences are then clustered based on their motif composition. It also offers the option to perform multiple sequence alignment based on motif.

### 4.1. Algorithm

#### 4.1.1. Motif discovery and annotation

MotifScope identifies repeat motifs across a given set of input sequences S = {s_1_, s_2_,…, s_n_} based on highly occurring k-mers (see Supplementary Algorithm 1 for implementation details). Due to the repetitive nature of TRs, a single k-mer *k*_*x*_ in a tandemly repeated sequence can yield a set of distinct k-mers (*K*_*x*_) as the repeat motif can be circularly permuted, for example for k-mer *k*_AAG_, *K*_AAG_ = {“AAG”, “AGA”, “GAA”}.

k-mer frequencies for varying lengths of k (the user can specify the maximum size of k-mer to screen) are first computed across all sequences. To that end, input sequences are concatenated into a string (*s*_*combined*_= ”{s_1_}${s_2_}$…${s_n_}”, and a suffix array and LCP array is computed for *s*_combined_ using the libsais library (available at https://github.com/IlyaGrebnov/libsais) (Nong et al. 2011). We iteratively walk through the suffix array, while keeping a set of active k-mers and their count. At each position *i* in the suffix array, the count of active k-mers with a *size ≤ LCP[i]* is increased by one. Active k-mers with a *size > LCP[i]* are stored in a result list, accompanied by the count with which they occurred in the sequence *s*_combinded_.

k-mers that can be represented as repetition of shorter sequences are excluded, e.g., “AGAG” is removed because it can be viewed as (AG)_2_. Also, k-mers are not considered if they contain the sequence separation symbol “$” or occur only once.

The final list of these k-mers are then sorted based on *k* × count so that k-mers that can mask longer sequences will be considered first. Given the set of sorted k-mers K = {k_a_, k_b_, k_c_,…}, MotifScope then iteratively identifies and annotates them across the input sequences (Algorithm 1) . In brief, for each iteration, a k-mer *k*_*j*_ is selected as the candidate motif, and the maximum continuous masked sequence length *l*_*mcs*_(*k*_*j*_) is determined. This is done for all k-mers, until *k* × count of the next k-mer is smaller than the maximum value of *l*_*mcs*_ that has already been observed for previously considered k-mers. The k-mer with the largest *l*_*mcs*_ value is subsequently selected, and masked from the sequence. The masking operation is detailed in Supplementary Algorithm 3.

The unmasked sequences are then used as the input for the subsequent iteration to discover the next candidate motif. For the i^th^ iteration (where i > 1), an additional step is taken to provide a canonicalized description of candidate motifs (Supplementary Algorithm 2). For example, if the set of already selected k-mers M = {“TGAGA”}, the next candidate motif, m_2_, is canonicalized towards “TGAGC” instead of one of its cyclical rotations “GAGCT”, “AGCTG”, “GCTGA” or “CTGAG”. This aims to ensure that these motifs are in a comparable representation, enabling clearer comparison between them. To achieve this, MotifScope considers all cyclical rotations of candidate k-mer m_i_, and selects the rotation that produces the maximum sum of pairwise alignment scores compared to all previously identified candidate motifs in M. This k-mer is then considered canonicalized and is added to M.

This k-mer selection process stops when the longest remaining sequence ≤ 1 bp in length or the length of the identified motif is 1 bp. All single occurrences of previously identified candidate motifs in the remaining sequences are then tagged with the corresponding motif as well. The bases that remain uncharacterized are then tagged with the single nucleotide at that position. In this way, each base in the sequences is assigned to one motif.

**Algorithm 1.**
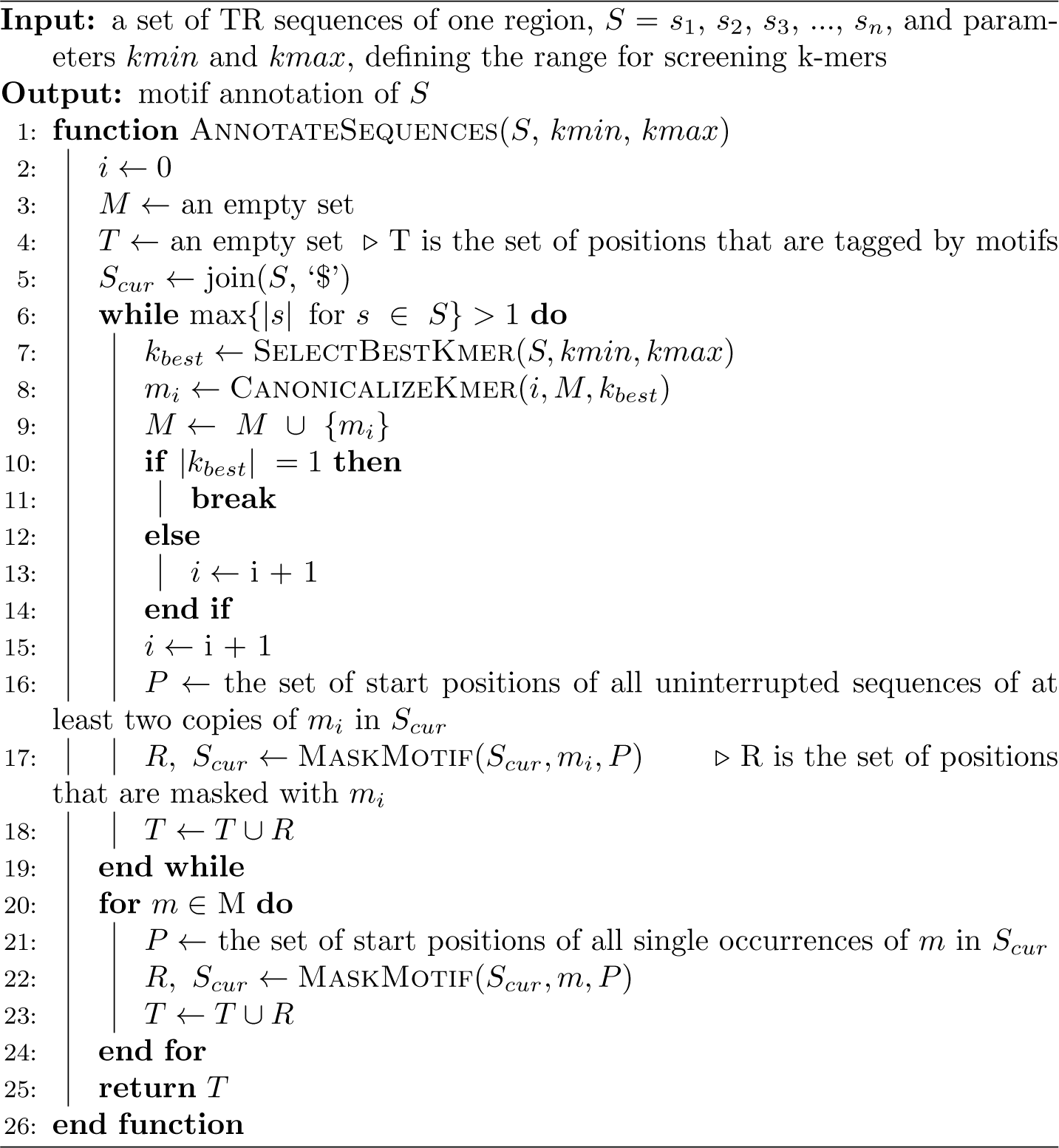
Motif annotation

#### 4.1.2. Clustering and alignment

To effectively compare the patterns of motifs in sequences and to enable further downstream analysis, MotifScope clusters sequences based on their motif organization and length. Each unique motif is assigned to a unique ASCII character, allowing for up to 248 unique motifs. Nucleotide sequences are then translated into a ‘motif sequence’ with these characters according to the motif assignments. Hierarchical clustering is performed on the pairwise edit distance matrix of these motif sequences.

MotifScope also includes the option for MSA using the mafft (v7.505) algorithm (Katoh and Standley 2013). MSA is done at the level of the motif sequences.

#### 4.1.3. Dimensional reduction of motifs to a color map

MotifScope accurately represents the underlying sequences, and uses a color-based visualization to display TR composition. It reflects motif similarity in a color spectrum such that similar color corresponds to similar motifs. This is achieved by projecting the pairwise alignment score matrix of motifs to a one-dimensional space by performing Uniform Manifold Approximation and Projection (UMAP).

### 4.2. Benchmarking

We benchmarked MotifScope’s ability to identify motifs and accurately represent TR sequences in the context of recently developed methods: uTR, vamos, and TRF. To do so, we used HG002 genome assembly, and a set of TRs from the PacBio repeat catalog (version 0.3.0, available at https://github.com/PacificBiosciences/trgt/tree/_main/repeats), which contains 171,146 TRs. One existing tool, vamos, can only be applied to locations in its repeat catalog, consisting of 467,104 VNTRs. The size distribution of TRs within the PacBio catalog differed from that in the vamos VNTR catalog; specifically, repeats were smaller in the PacBio catalog, with 99.97% of repeats being ≤ 100 bp in the GRCh38 human reference genome, while in the vamos VNTR catalog, 80.42% of repeats are ≤ 100 bp (Figure 10A). The PacBio catalog also showed reduced sequence complexity, as illustrated by the distinct k-mer (k = 10) counts observed in each repeat in the GRCh38 human reference genome (Figure 10B).

**Figure 10.**
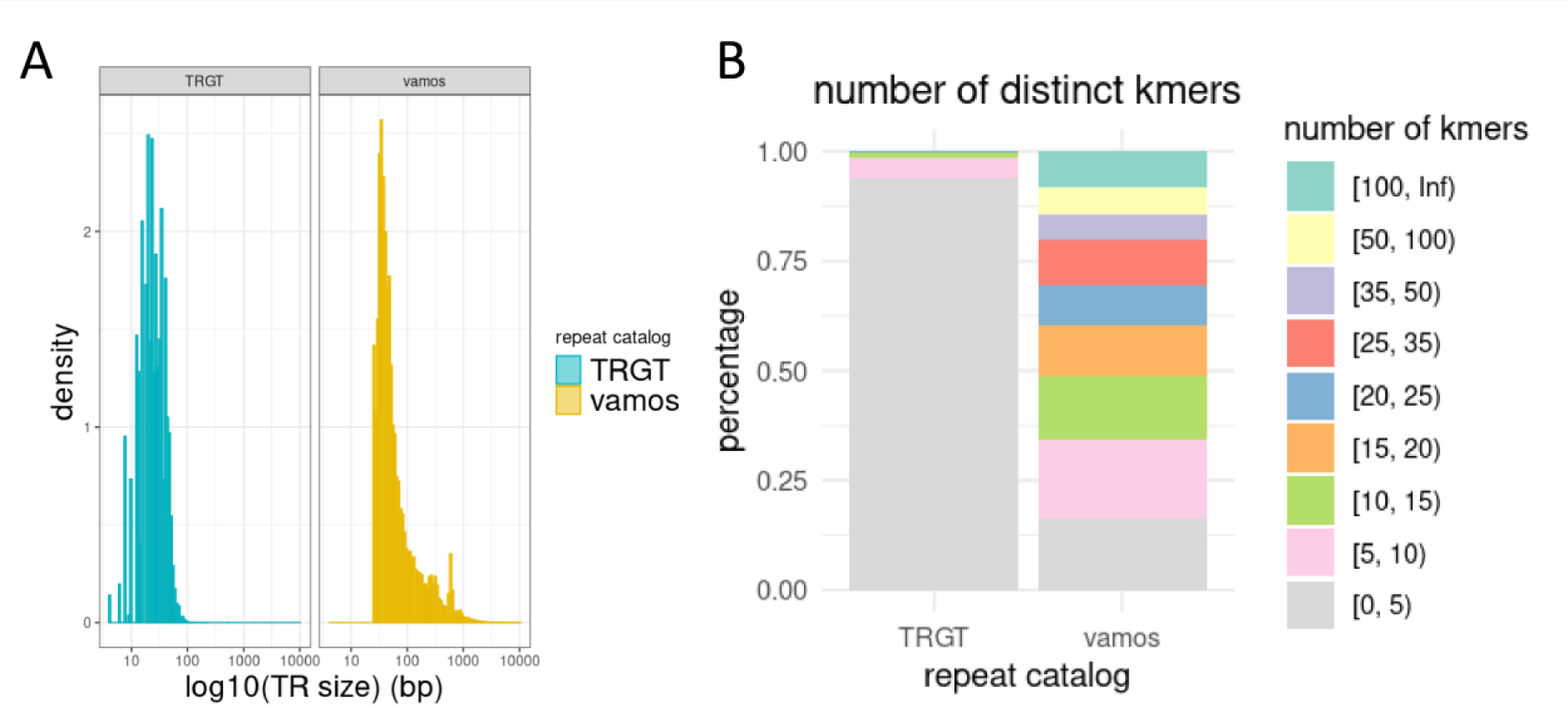
Summary of the PacBio TRGT repeat catalog and the vamos VNTR catalog. The figure provides an overview of two catalogs: the PacBio TRGT repeat catalog, comprising 171,146 repeats, and the vamos VNTR catalog, comprising 467,104 repeats. (A) compares the size distribution of these two datasets. (B) presents a stacked bar plot showing the number of distinct kmers (k=10) for each locus in both datasets in GRCh38.

We evaluated the tools performances based on the overlap between these two catalogs, which constituted 5,486 VNTRs. Additionally, to provide a comprehensive evaluation, we also randomly sampled 5,000 repeats from the vamos VNTR catalog and compared the performances between these four tools on this set of repeats.

For vamos, several motif databases have been made available by the authors: “vamos original”, which uses motifs identified in samples from the HPRC and HGSVC; and “vamos efficient”, in which rare motifs have been replaced with more common ones while ensuring a bounded total replacement cost (compression strength q = 0.2) (Ren et al. 2023).

### 4.3. Sequencing data

#### Public sequencing data

Individual PacBio HiFi reads as well as whole genome assemblies of the paternal and maternal haplotypes of HG002 genome were used for benchmarking (Wang et al. 2022). To assess repeats across different individuals, whole genome assemblies of the HPRC samples were also used (Wang et al. 2022).

#### CANVAS patients

The PacBio HiFi sequencing data of a blood sample of a Dutch CANVAS patient was additionally used to evaluate the performance of different tools in a clinical setting based on the RFC1 repeat (van de Pol et al. 2023). The genome of the Dutch CANVAS patient was assembled with hifiasm (version 0.16). In addition, PacBio HiFi sequencing data of a blood sample of another Dutch CANVAS patient was used to show read variability (van de Pol et al. 2023).

#### Dutch AD patients and cognitively healthy centenarians

PacBio HiFi sequencing data of 250 AD patients and 238 Dutch cognitively healthy centenarians were additionally used for multi-genome comparisons of TR motifs. Additional information about the sequencing and data processing can be found in Salazar et al., 2023 (Salazar et al. 2023). Targeted local assembly of the regions of interest was done on these genomes using Otter on HiFi reads (available at https://github.com/holstegelab/otter).

### Data access

MotifScope is available as an open-source tool on GitHub at (https://github.com/holstegelab/MotifScope)

Whole genome PacBio HiFi sequencing data from the HPRC is available from https://github.com/human-pangenomics/HPP_Year1_Data_Freeze_v1.0.

The PacBio repeat catalog is available from https://github.com/PacificBiosciences/trgt/blob/main/repeats/repeat_catalog.hg38.bed.

The vamos VNTR catalog, both the original motif set, and an efficient motif set at the level of Delta=20, can be accessed from the vamos GitHub repository at: https://github.com/ChaissonLab/vamos.

The sequencing data of the Dutch AD patients and centenarians used in the present study can be made available for analysis upon request to the authors.

## Author contributions

Conceived the study: HH; Wrote the manuscript: YZ, AS, MH, NT, HH; Patient selection: NT, HH, E-JK, Patient blood collection and sequencing: SW, JK LK, E-JK; Data management: NT, MH, SvdL, Bioinformatic analysis: YZ, MH.

## Supporting information

Supplementary Algorithm 1

Supplementary Algorithm 2

Supplementary Algorithm 3

## Acknowledgements

The authors are grateful to all study participants, their family members, the participating medical staff, general practitioners, pharmacists and all laboratory personnel involved in patient diagnosis, blood collection, blood biobanking, DNA preparation and sequencing. Part of the work in this manuscript was carried out on the Cartesius supercomputer, which is embedded in the Dutch national e-infrastructure with the support of SURF Cooperative. Computing hours were granted to H. H. by the Dutch Research Council (‘100plus’: project# vuh15226, 15318, 17232, and 2020.030; ‘Role of VNTRs in AD’; project# 2022.31, ‘Alzheimer’s Genetics Hub’ project# 2022.38). This work is supported by a VIDI grant from the Dutch Scientific Counsel (#NWO 09150172010083) and a public-private partnership with TU Delft and PacBIo, receiving funding from ZonMW and Health∼Holland, Topsector Life Sciences & Health (PPP-allowance), and by Alzheimer Nederland WE.03-2018-07. H.H., S.L., are recipients of ABOARD, a public-private partnership receiving funding from ZonMW (#73305095007) and Health∼Holland, Topsector Life Sciences & Health (PPP-allowance; #LSHM20106). S.L. is recipient of ZonMW funding (#733050512). H.H. was supported by the Hans und Ilse Breuer Stiftung (2020), Dioraphte 16020404 (2014) and the HorstingStuit Foundation (2018). Acquisition of the PacBio Sequel II long read sequencing machine was supported by the ADORE Foundation (2022).

HH has a collaboration contract with Muna Therapeutics, PacBio, Neurimmune and Alchemab. She serves in the scientific advisory boards of Muna Therapeutics and is an external advisor for Retromer Therapeutics. Research of Alzheimer center Amsterdam is part of the neurodegeneration research program of Amsterdam Neuroscience. Alzheimer Center Amsterdam is supported by Stichting Alzheimer Nederland and Stichting Steun Alzheimercentrum Amsterdam. The clinical database structure was developed with funding from Stichting Dioraphte.

