## Supplementary Algorithm 1 for "MotifScope: a multi-sample motif discovery and visualization tool for tandem repeats"

Input: a set of TR sequences of one region, S, and parameters kmin and kmax, defining the range for screening kmers.

Output: the best kmer to annotate the sequences $k_{best}$

```
Function SelectBestKmer(S, kmin, kmax)
    Kmer_Counts <- an empty set
    for k <- kmin to kmax do
        kmers <- a set contains all kmers present in S
        foreach kmer in kmers
            Kmer_Counts <- Kmer_Counts ∪ {(kmer, count, k*count)}
    Sorted_Kmers <- Kmer_counts sorted based on k*count in descending order
    max_mask <- 0
    l_mcs <- an empty set
    foreach (kmer, count, k*count) in Sorted_Kmers do
            if k*count >  max_mask then
                l_mcs(kmer)<- the longest length of uninterrupted sequence with kmer (≥ 2 copies of kmer) among S
                max_mask = max(l_mcs(kmer), max(l_mcs))
            else
                break
    k_best <- kmer with max_mask
    return k_best
```
