## Supplementary Algorithm 2 for "MotifScope: a multi-sample motif discovery and visualization tool for tandem repeats"

Input: the iteration number i, the set of previously identified motifs M, the best kmer, kbest, for annotation in this iteration

Output: canonicalized kmer as the candidate motif of this iteration mi

```
Function CanonicalizeKmer(i, M, k_best)
    if i == 1 then #if it is the first iteration
        m_i <- k_best
    else
        K_best <- {k_best[i:] + k_best[:i] for i <- 0 to k}
        Kmer_Alignment_Score <- an empty set of (kmer, Sum_Alignment_Score)
        foreach kmer in K_best do
            Sum_Alignment_Score <- 0
            foreach m in M do
                Alignment_Score <- pairwise alignment score between kmer and m
                Sum_Alignment_Score <- Sum_Alignment_Score + Alignment_Score
            Kmer_Alignment_Score <- Kmer_Alignment_Score ∪ {(kmer, Sum_Alignment_Score)}
        Sorted_Alignment <- Kmer_Alignment_Score sorted based on Kmer_Alignment_Score in descending order
        m_i <- Sorted_Alignment[0]
    return m_i
```
