## Supplementary Algorithm 3 for "MotifScope: a multi-sample motif discovery and visualization tool for tandem repeats"

Input: a single TR sequence $S_j$, motif m to annotate the sequences, a set contains all start positions of m in s to be masked, Start_Positions

Output: a set of regions that are annotated with m and the remaining sequences

```
Function MaskMotif(Sj, m, P)
    Masked_Regions <- an empty set
    foreach position in Start_Positions do
    Masked_Regions <- Masked_Regions ∪ {(j, m, position, position + |m|)}
    foreach (j, m, position, position + |m|) in Mask_Regions
        S <- S - Sj
        Sj <- the remaining sequence of Sj after masking [position, position + |m|] in Sj
        S <- S ∪ {Sj}
    return Masked_Regions, S
```
