## Supplementary Figure 1 for "MotifScope: a multi-sample motif discovery and visualization tool for tandem repeats"

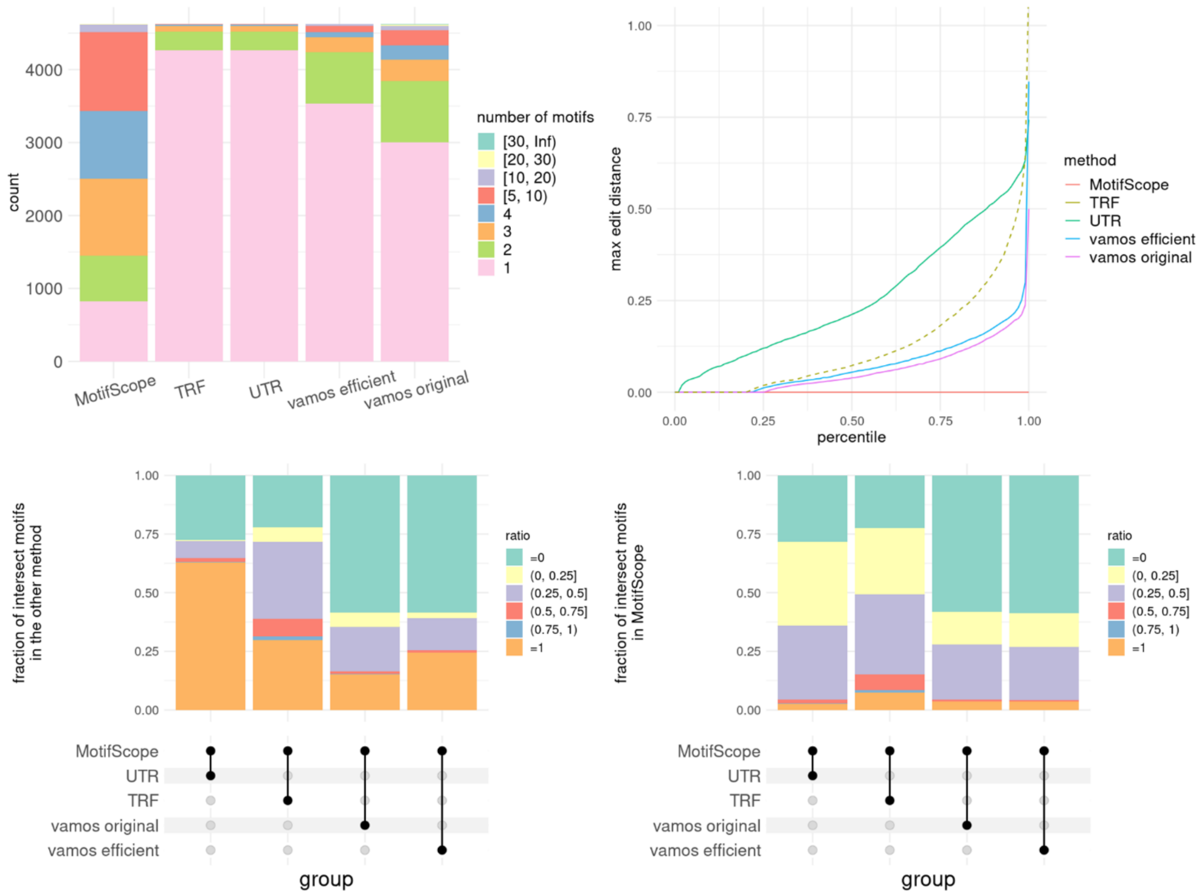


**Supplementary Figure 1. Comparative analysis of tandem repeat characterization in vamos repeat catalog. Randomly sampled 5000 regions.** Four tools were tested in the analysis, MotifScope, TRF, uTR, and vamos. For vamos, here we used both the original motif set (vamos original) and the efficient motif set (vamos efficient). (A) shows the number of motifs discovered by each of the four tools. (B) shows the normalized edit distance between the actual sequence and the results obtained from the four tools, normalized with respect to repeat length. (C) and (D) shows the intersection of motifs between MotifScope and other tools (dots connected by lines below the X axis): for (C), the stacked bar plot at the top shows the fraction of intersected motifs over the total number of motifs found by the other tool; for (D) the stacked bar plot at the top shows the fraction of intersected motifs over the total number of motifs found by MotifScope.
