## Supplementary Figure 2 for "MotifScope: a multi-sample motif discovery and visualization tool for tandem repeats"

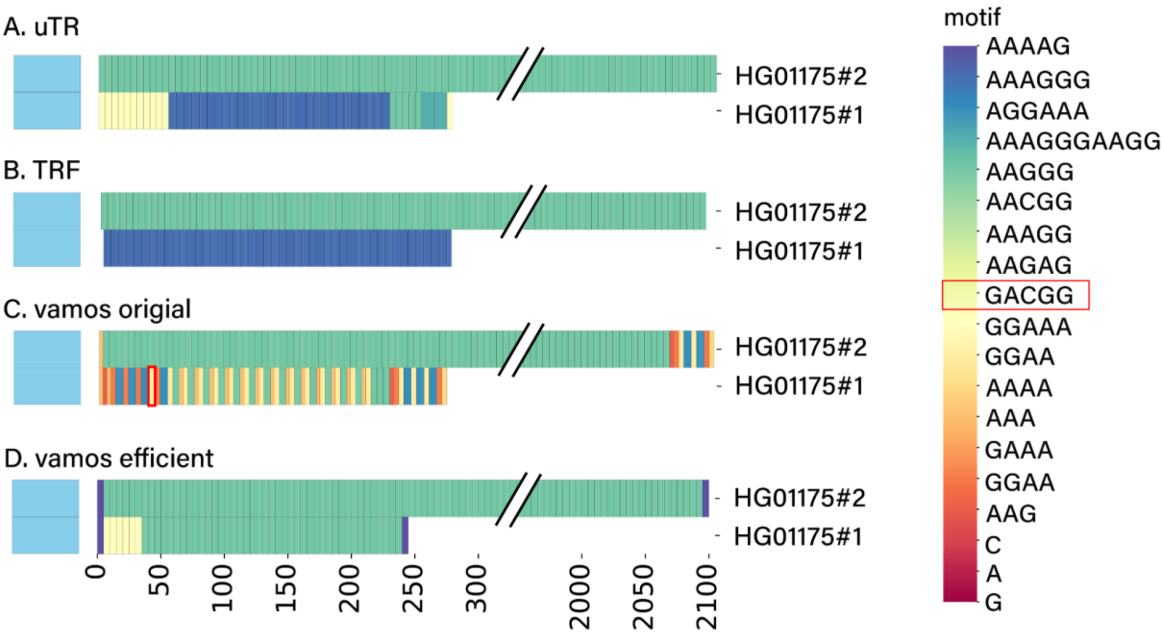


**Supplementary Figure 2**. **Motif characterization of the RFC1 repeat in HG01175 with different tools.** The results from (A) uTR, (B) TRF, (C) vamos original and (D) vamos efficient of the RFC1 repeat in the HG01175 assembly. The “GACGG” motif is highlighted in a red box.
