## Supplementary Figure 3 for "MotifScope: a multi-sample motif discovery and visualization tool for tandem repeats"

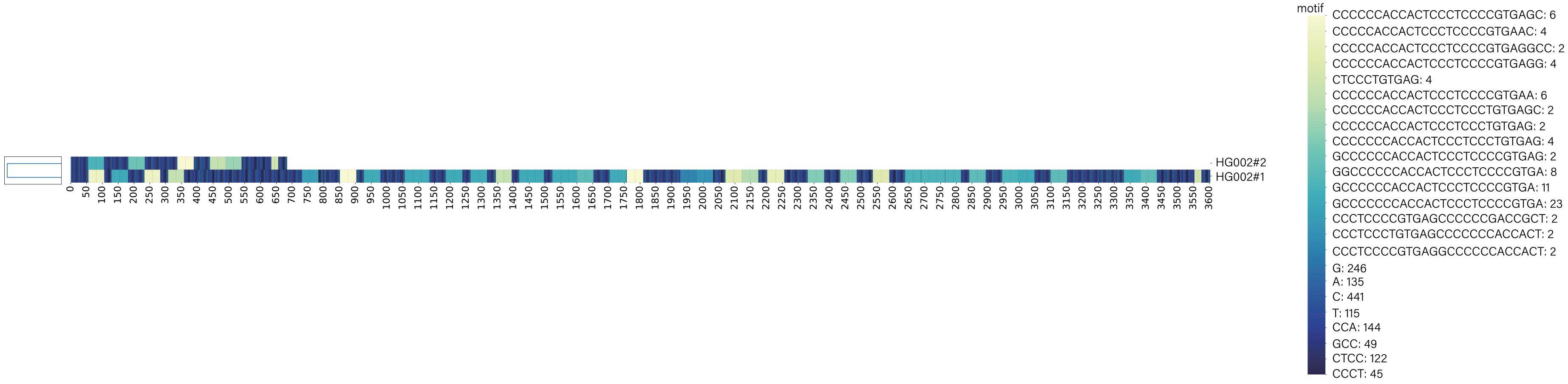


**Supplementary Figure 3. Motif characterization of the ABCA7 repeat in HG002 with MotifScope.** This shows the visualization of the decomposing result from MotifScope of the local assembly of the ABCA7 repeat in HG002 genome assembly, with 10 bp flanking on both sides. Distinct motifs are represented in different colors, as indicated by the color bar on the right side of the figure, followed by the occurrence of motifs in the figure.
